# Antibody targeting the anti-parallel topology of human telomeric G-quadruplex DNA

**DOI:** 10.1101/2025.05.13.653741

**Authors:** Nasab Reda, Sofia Muret, Clément Estève, Emilie Derathe, Maria Fidélia Susanto, Eve Pitot, Hugues Bonnet, Thomas Lavergne, Dennis Gomez, Jérôme Dejeu, Natale Scaramozzino, Eric Defrancq

## Abstract

G-quadruplexes (G4s) are four-stranded nucleic acid structures that have gathered a significant attention due to their involvement in key biological processes, including gene regulation, genome stability, and telomeres maintenance. Some G4 antibodies have been developed to selectively recognize these structures over duplex DNA; however, most, even the widely studied BG4 and 1H6, bind G4s in a general manner and lack discrimination between distinct topologies, particularly between parallel and antiparallel conformations. In this study, we report on the development and characterization of a novel antibody selected via phage display method using a constrained antiparallel G4 structure mimicking one of the conformation adopted *in vitro* by the human telomeric sequence. Our findings demonstrate that this new antibody selectively recognizes the antiparallel topology of the telomeric G4 sequence, a property further validated in cellular models.

## Introduction

Over the past two decades, a large body of research has revealed that genomic DNA can form alternative structures beyond the well-known canonical double helix (B-form antiparallel right-handed double helix). These non-B DNA structures can significantly impact both genetic and epigenetic processes. Among these, G-quadruplexes (G4s) have attracted considerable attention due to their potential biological relevance. G4s are tetrahelix structures that arise in G-rich sequences and result from the self-assembly of guanine moieties into G-quartets, which are further stabilized by π-stacking interactions and coordination with K^+^ or Na^+^ metal cations. ^1^ They are found in critical genomic regions including telomeres, promoters, and untranslated regions. In particular, they play crucial roles in various cellular processes, including the regulation of gene expression, the maintain of genome integrity, the control of chromosomal stability and telomere maintenance.^2^ They have also found significant interest as potential therapeutic targets. In cancer research, for instance, the presence of G4 structures at telomeric regions has been linked to the inhibition of telomerase activity, a key mechanism underlying the unlimited proliferation of cancer cells.^3^

To investigate the structural and functional properties of G4 DNA, numerous chemical and biological tools have been developed. A diverse array of small molecules, often referred as G4 ligands, have been described for their ability to specifically bind and stabilize G4 structures.^4, 5, 6, 7^ These ligands are invaluable tools for probing the existence and biological relevance of G4s. In parallel, antibodies have emerged as powerful biological tools to explore the functional roles of G4 DNA *in vivo*. A limited number of G4-targeting antibodies, including 1H6 reported by Lansdorp and Coll.^8^ and BG4 from Balasubramanian and Coll., ^9^ have been developed and selected for their ability to selectively recognize G-quadruplexes over duplex DNA. The BG4 antibody stands out as the most well-characterized and widely used tools for G4 detection. BG4 preferentially binds to G-quadruplexes, providing a valuable means for studying G4 biology *in vitro* and *in vivo*.

A defining feature of G4s is their intrinsic polymorphism. Numerous *in vitro* studies have shown that G4s are highly susceptible to adopt multiple topologies, which exist in dynamic equilibrium. The specific topology adopted by a G4 structure depends on several factors, including the length and sequence composition of the forming strands, as well as environmental conditions such as the concentration and nature of metal cations, and the local molecular crowding environment. ^10^ For example, the telomeric sequence d[AGGG(TTAGGG)_3_] can form different G4 topologies, including parallel, antiparallel, or hybrid configurations, depending on the conditions.^11^

This structural polymorphism poses a significant challenge in the study of G4 recognition by biological components and complicates global studies of structure-activity relationships of G4-interacting proteins. To date, only a limited number of G4 antibodies have been identified that can selectively bind G-quadruplexes. Most of them, including BG4 and 1H6, recognize the G4 structure in a general sense but are not specific to a particular topology, especially when it comes to differentiate between parallel and antiparallel G4 conformations.^8, 9^

In an effort to overcome the limitations due to the G4 structural heterogeneity, we have developed a strategy to constrain the accessible topologies of G-quadruplex-forming sequences to a single, well-defined structure. This innovative approach, called Template-Assembled Synthetic Quadruplex (TASQ), utilizes a rigid cyclic peptide scaffold to direct the intramolecular assembly of anchored oligonucleotides into a specific G-quadruplex topology. By using this strategy, we have successfully prepared both fully parallel tetrameric DNA-G4 **1** and an antiparallel G-quadruplex **2** (Fig. 1), both mimicking the respective conformations adopted *in vitro* by the human telomeric sequence.^12,13^

**Figure 1:**
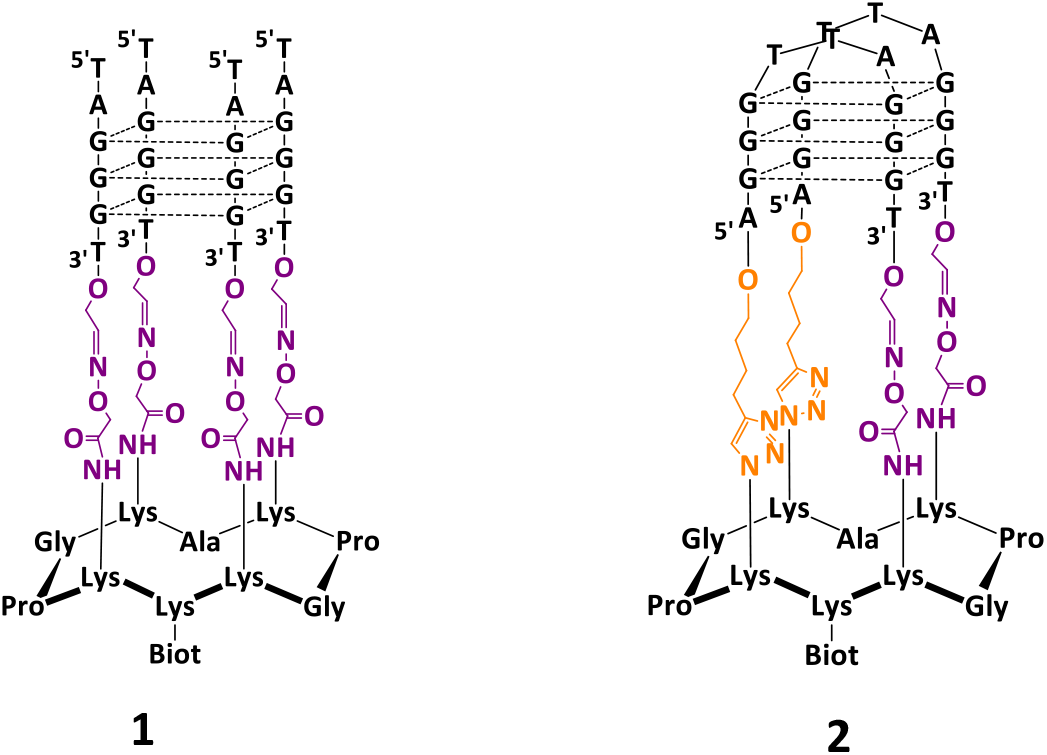
Structure of constrained G-quadruplex DNA using TASQ concept mimicking parallel **1** and antiparallel **2** topology of the telomeric sequence.^12, 13^

Both systems **1** and **2** exhibit exceptional stability, with melting temperature (*ΔTm*) increases of over 20°C compared to unconstrained G4-DNA, and they served as robust models for studying ligand binding properties in a controlled and reproducible manner.^14^ These G4 constrained structures also offer an opportunity to study the recognition of specific G4 topologies by proteins, ^15^ and in the present case, could serve as a powerful tool for developing antibodies capable of distinguishing between different G4 topologies.

Building on these unique characteristics we used the system **2** as the target for the selection of an antibody that selectively recognizes an antiparallel G4 topology while system **1** served as control of parallel topology. Such antibody would represent an invaluable tool for investigating the biological roles of the antiparallel topology of G4 telomeric sequence.

## Results and discussion

### Selection of scFv **DST20** by phage display

To select the antibody candidates as single chain fragment variable format (*scFv*), we employed the phage display technique with modified biopanning procedures that enabled the identification of **DST20** (see Methods in Supporting Information). Phage display is a method that presents peptides on the surface of filamentous bacteriophages and is the most effective technique for generating antibodies through selection and amplification cycles.^16^ In this study, we used the bacteriophage M13, where a *scFv* fragment was fused to the PIII protein, an envelope protein expressed in 3-5 copies. A large diversity of *scFv* fragments was generated from the human *Tomlinson* library, ^17^ which contains approximately 10^8^ different *scFv* sequences. The phagemid pIT2, carrying the *scFv* sequence displayed on the phage surface, was transmitted to bacteria (TG1) following infection.

The selection cycles were carried out on the telomeric G4 DNA antiparallel topology **2**, (Fig. 1) which allowed to preserve the antiparallel topology through the phage display experiments. The selection strategy consisted of presenting the target **2**, which was attached to streptavidin beads (thanks to the biotin present on **2**), to the phage library (*i*.*e*. human *Tomlinson* library). To identify the *scFvs* from the phage display selection cycles, enzyme-linked immunosorbent assay (ELISA) tests were carried out on approximately one hundred isolated candidates. These tests were performed on the supernatants of the bacteria media infected with each phage, following induction by IPTG, releasing *scFv* fused to the PIII protein of phage M13 as described in the protocol. The amount of generated *scFv* varies greatly from one candidate to another, so we ranked the candidates in relation to the ELISA results according to their response to the G4 telomeric target **2** *versus* duplex DNA (HP-DNA in Table 1). Candidate **DST20** was selected because it gave the best responses, *i*.*e*. a positive ELISA signal for target **2** and no signal for HP-DNA (See the Supporting Information, Fig. S1).

**Table 1.**
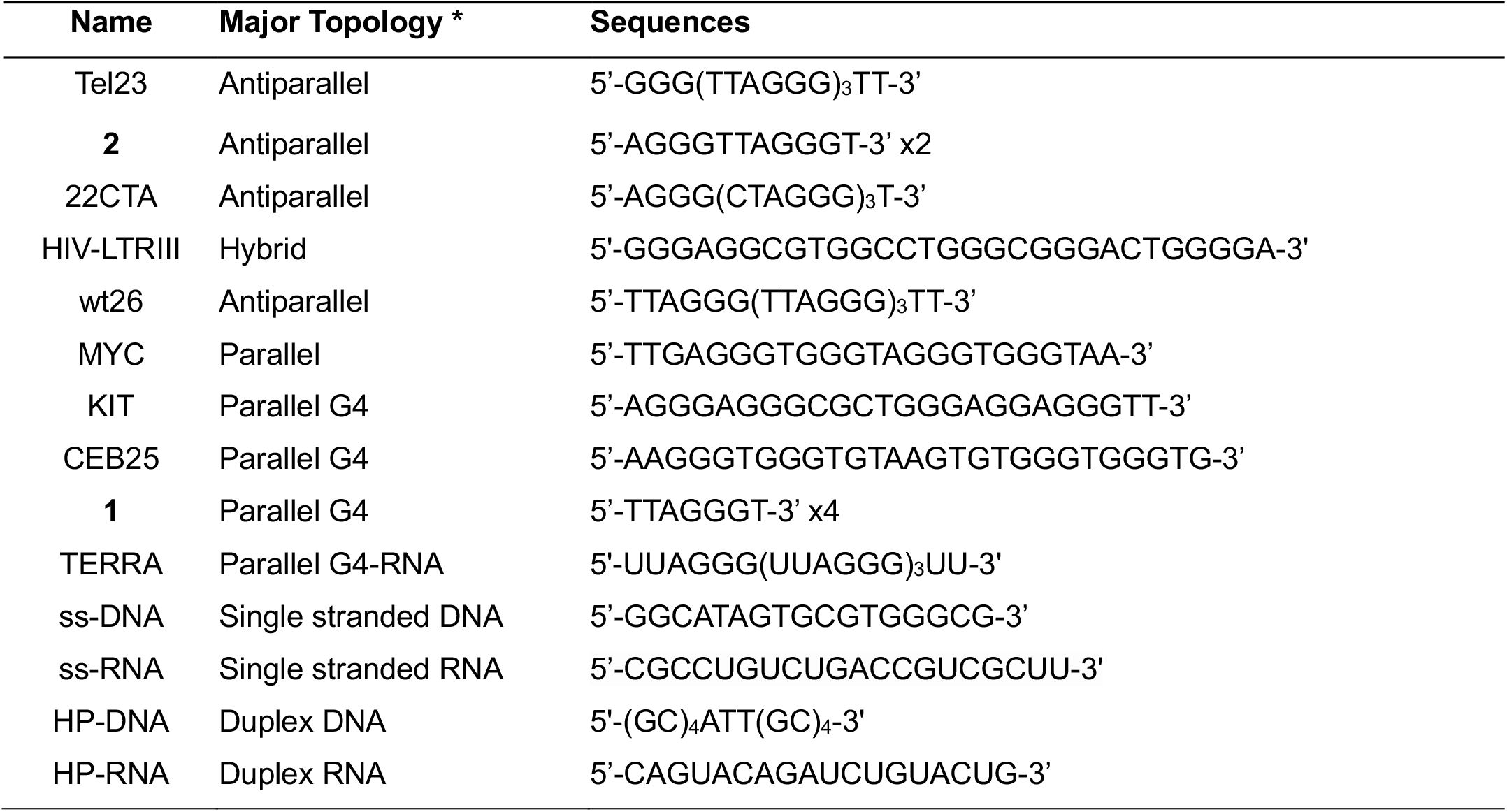
Sequences used to study the selectivity of **DST20** antibody. **\*** The topology was determined by CD analysis in the conditions used for the phage display selection (PBS buffer at pH 7.2). See Figures S4-S6 in the Supporting Information for CD spectra.

Phagemid expressing the *scFv* **DST20** were then sequenced to determine the amino acid sequences of both the framework and variable regions. **DST20** was then produced in CHO cells culture as *scFv* and *IgG* formats for further characterization.

### Extensive ELISA tests to verify the selectivity of the **DST20** antibody

The selectivity of the **DST20** antibody in both formats (*scFv* and *IgG*) for the antiparallel telomeric G4 DNA over other nucleic acid structures was first evaluated using ELISA. A diverse set of oligonucleotide sequences, adopting antiparallel, hybrid, parallel G4 structures (as determined by CD experiments in phage display conditions), as well as duplex DNA and RNA and both single-stranded DNA and RNA, was tested (Table 1).

*IgG* **DST20** exhibits a strong affinity for the antiparallel G4 structures formed by the telomeric sequences (found in the constrained G4 DNA **2** and Tel-23 in these conditions) and for wt26, which consists of a mixture of topologies including major antiparallel (Fig. 2a and 2b). In contrast, no binding is observed with the parallel DNA G4 sequences (MYC, KIT, CEB25 and constrained system **1**) or with the parallel TERRA RNA sequence even at *IgG* **DST20** concentrations up to 200 nM (Fig. 2c and 2d). Similarly, no interaction is detected with double-helix DNA or RNA (HP-DNA and HP-RNA, Fig. 2e). A weak interaction is observed with the single stranded DNA model (Fig. 2f) whereas no binding is detected with the single-stranded RNA model. Interestingly, no interaction is detected between *IgG* **DST20** and the antiparallel G4 22CTA, a mutant of the telomeric sequence (Figure 2a), despite only a single nucleotide variation in the loop compared to the telomeric sequences Tel-23 and wt26, which contain TTA instead of CTA.

**Figure 2:**
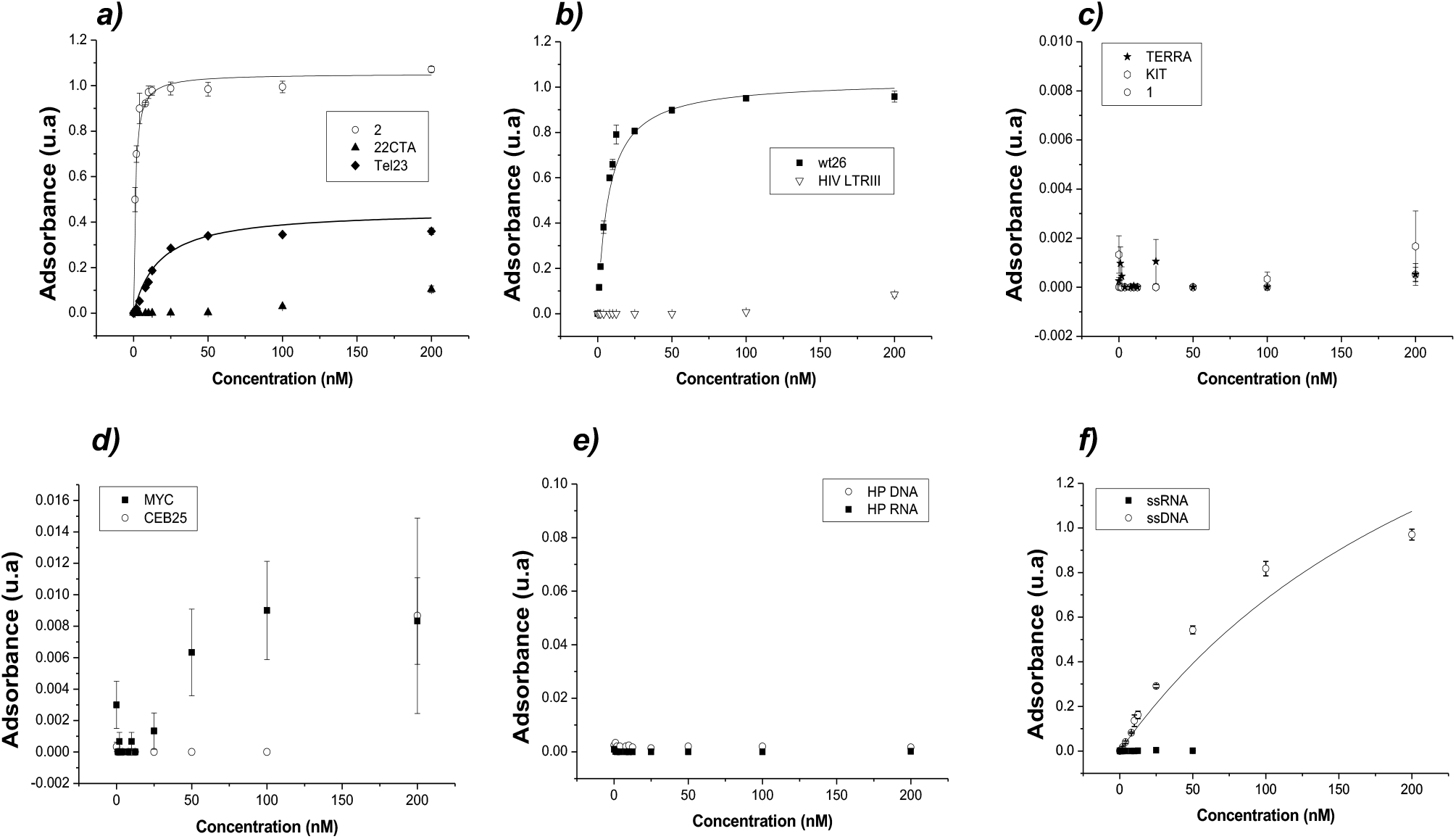
Structure specificity of the *IgG* **DST20** antibody for G4 structures. Binding curves as determined by ELISA with ***a)*** the antiparallel G4 Tel23, 22CTA and constrained G4 **2** targets, ***b)*** the mixture of parallel and antiparallel topologies wt26 and hybrid HIV LTRIII targets, ***c)*** the parallel G4 TERRA, KIT and constrained structure **1, *d)*** the parallel G4 MYC and CEB25, ***e)*** the double stranded DNA and RNA models and ***f)*** the single stranded DNA and RNA models.

ELISA assay was also performed with BG4 antibody (*IgG* format) for comparison. An approximative *K*_*D*_ value for the interaction with Tel23 DNA G-quadruplex of 17 nM was obtained which is similar to that measured for *IgG* **DST20** (Figure S2).

The *scFv* **DST20** exhibited a binding profile similar to that of *IgG* **DST20**: demonstrating good selectivity for antiparallel G4 DNA over parallel G4 DNA as well as over duplex and single stranded DNA and RNA (See Figure S3 in the Supporting Information). However, the affinity of *scFv* **DST20** for the antiparallel DNA target **2** (approximate *K*_*D*_ = 100 nM from Figure S3) was significantly lower – by a factor of 100 - compared to *IgG* **DST20** (approximate *K*_*D*_ = 1 nM for **2**). ^18^ Based on these results, the selectivity of **DST20**, specifically in its *IgG* format, was further investigated using Bio-Layer Interferometry (BLI) with a panel of diverse sequences (Table 1).

### Confirmation of selectivity of **DST20** antibody by BioLayer Interferometry

The BioLayer Interferometry (BLI) was used to study the interaction of *IgG* **DST20** with various G4 and non-G4 forming sequences (Table 1). In a first experiment, the different 3’-biotinylated RNA or DNA sequences as well as the constrained systems **1** and **2** were immobilized on the streptavidin sensor before dipping them into 2 nM solutions of *IgG* **DST20** (sensorgrams in Fig. S7) for 20 minutes. Figure 3 shows the normalized signals for each sequence.

**Figure 3:**
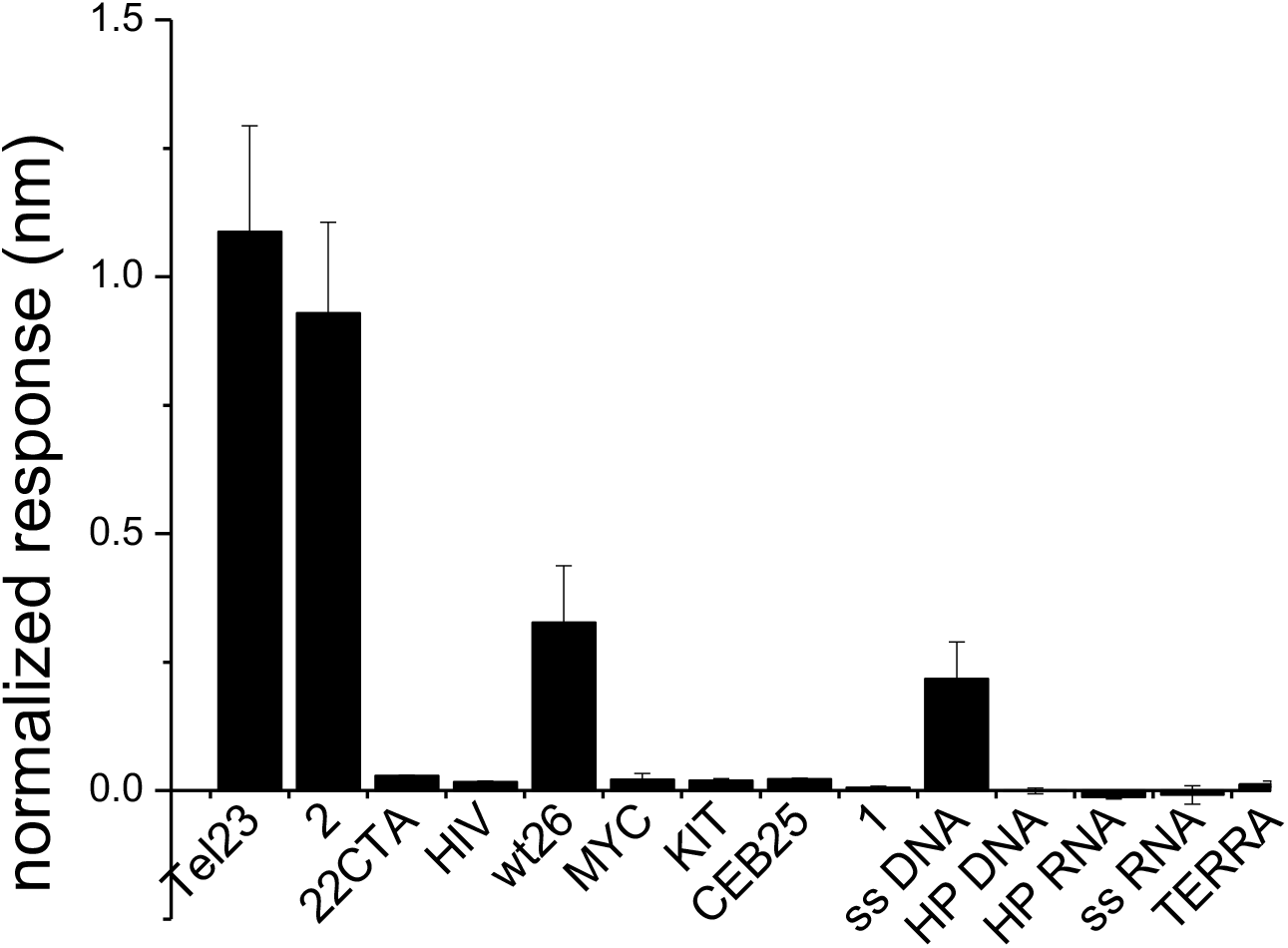
Normalized response obtained for the binding of *IgG* **DST20** at 2 nM with different DNA and RNA sequences, showing that *IgG* **DST20** has an affinity for antiparallel G4 sequences Tel23, **2** as well as for wt26 and ss-DNA and negligible binding with parallel G4 sequences MYC, KIT, CEB25, **1** and duplex DNA and RNA (HP-DNA and HP-RNA) as well as with ss-RNA. The normalized response consists in the ratio of the observed signal to the background signal (*i*.*e*. immobilization signal).

The normalized responses confirm the results obtained from ELISA assays (Fig. 3). *IgG* **DST20** binds the sequences forming the antiparallel form of the telomeric DNA Tel23, **2** as well as with wt26, and a slight interaction is detected with ss-DNA and negligible binding with all the parallel, hybrid and alternative antiparallel (*i*.*e*. not telomeric) forming G4 sequences as well as with double stranded DNA is observed. However, using only the intensity of the BLI signals as a measurement of binding can lead to biases, since the intensity of BLI signals is known to strongly depend on the size and conformation of the interacting molecules, both the one attached on the surface on the sensor and the dissolved analyte.^19,20^ Thus, the observations from the normalized responses were confirmed by the measurements of the affinity between the two partners (*i*.*e*. the antibody with the target DNA).

### Determination of the affinity constants

An approximative affinity was determined by ELISA (*vide supra*) and was used to adjust the experimental conditions for the precise measurement of the affinity by BLI. The different sensorgrams were presented in Figure S8 (Supporting Information). The shape of the curve depends on the sequences. The dissociation constant, *K*_*D*_, was determined from the Langmuir equation (response at the equilibrium *versus* the concentration, *i*.*e. K*_*D*_ *thermodynamic*) and also from the association (*k*_*on*_) and the dissociation (*k*_*off*_) kinetic constants (ratio *K*_*off*_*/k*_*on*_ *i*.*e. K*_*D*_ *kinetic*). The latter one was only determined for the G4 sequences (Table 2).

**Table 2:**
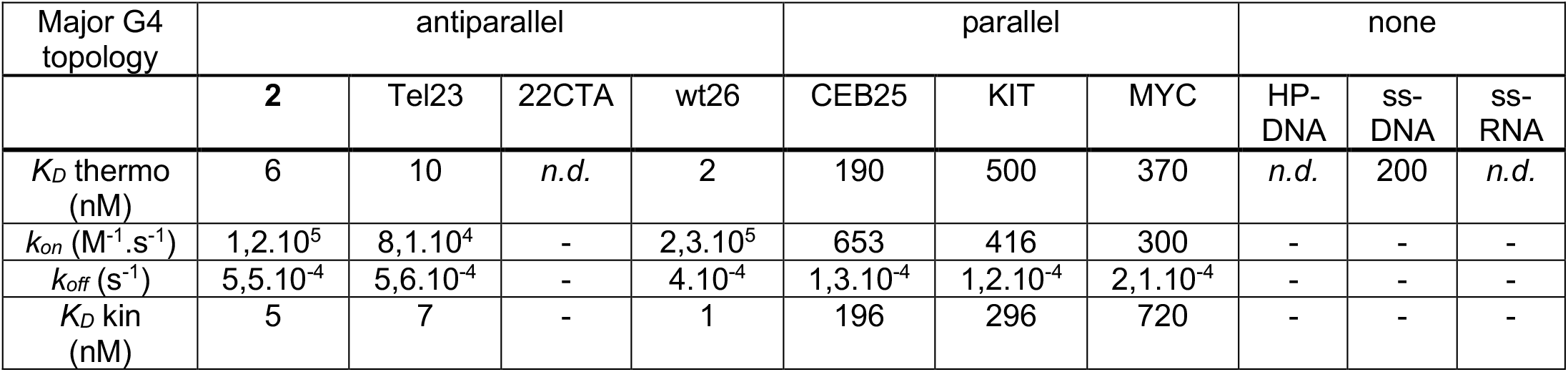
kinetic and dissociation constants for *IgG* **DST20** *versus* different DNA and RNA sequences. The concentration ranges from 2.5 nM to 1000 nM depending on the sequence (see SI). *n*.*d*.: the shape of the sensorgramm does not allow the exact determination of *K*_*D*_ value at the studied concentrations which indicates that the approximative *K*_*D*_ value is superior to 200 nM.

| Major G4 topology | antiparallel |  |  |  | parallel |  |  | none |  |  |
| --- | --- | --- | --- | --- | --- | --- | --- | --- | --- | --- |
|  | **2** | Tel23 | 22CTA | wt26 | CEB25 | KIT | MYC | HP-DNA | ss-DNA | ss-RNA |
| $K_D$ thermo (nM) | 6 | 10 | *n.d.* | 2 | 190 | 500 | 370 | *n.d.* | 200 | *n.d.* |
| $k_{on}$ ($M^{-1}.s^{-1}$) | $1,2.10^5$ | $8,1.10^4$ | - | $2,3.10^5$ | 653 | 416 | 300 | - | - | - |
| $k_{off}$ ($s^{-1}$) | $5,5.10^{-4}$ | $5,6.10^{-4}$ | - | $4.10^{-4}$ | $1,3.10^{-4}$ | $1,2.10^{-4}$ | $2,1.10^{-4}$ | - | - | - |
| $K_D$ kin (nM) | 5 | 7 | - | 1 | 196 | 296 | 720 | - | - | - |

A high affinity of *IgG* **DST20** was found for the antiparallel topology from the telomeric sequence (Tel23, wt26 and **2**) with a *K*_*D*_ value around 10 nM and a good concordance between thermodynamic and kinetic *K*_*D*_ values (Table 2). To confirm the selectivity of *IgG* **DST20** toward the antiparallel telomeric G4 sequence, the affinity for parallel G4 sequences (CEB25, KIT, MYC) was determined despite the very low signals observed during the screening experiment (Figure S7). An affinity was measured, although it was found to be more than 20 times weaker than that of the antiparallel sequences **2** and Tel23. The difference was due to a lower kinetic association constant (500 M^-1^.s^-1^ *versus* 10^5^ M^-1^.s^-1^ for the antiparallel G4) which also explained the weak amplitude of the signal observed during the screening (Figure 3).

The affinity for double-stranded and single-stranded DNA and RNA was also evaluated by BLI. A slight affinity for ss-DNA was measured with a *K*_*D*_ value around 200 nM. For the other structures, the shape or the absence of signal up to 500 nM concentration does not allow for a determination of the affinity constant whatever the method used suggesting a very weak interaction with those DNA and RNA structures.

Again, the BLI analysis confirms the absence of interactions of *IgG* **DST20** for the antiparallel 22CTA sequence. This result suggests that a structural feature of the telomeric sequence (*e*.*g*. the nucleotide composition of the loops) is decisive for the recognition by the antibody. The *K*_*D*_ measurements by BLI therefore confirm the high specificity of *IgG* **DST20** to bind the antiparallel G4 topology adopted by the telomeric sequence over parallel, hybrid and alternative antiparallel topologies of G-quadruplexes.

The stability of *IgG* **DST20** was also studied by SDS-PAGE, SEC-HPLC and BLI. *IgG* **DST20** was found to be stable after 1 month at r.t and 37°C with less than 5% of degradation.^21^ Measurements by BLI confirms this stability with a *K*_*D*_ value for the interaction with Tel23 remaining consistent over this time period.

### Visualization of G4 DNA structures in human cells

To explore the interaction of *IgG* **DST20** with G-quadruplex (G4) DNA structures in cellular environment, we performed immunofluorescence assays in U2OS and HeLa cells. Staining with the *IgG* **DST20** antibody revealed a distinct punctate nuclear pattern which was not observed in control samples lacking the antibody (Fig. S.10). In U2OS cells, approximately 80 **DST20**-positive foci *per* nucleus were detected, indicating a recognition of a limited set of genomic regions. Next, we investigated whether the *IgG* **DST20** antibody specifically targets telomeric G4 sequences rather than other G-quadruplex structures by examining its colocalization with TRF2, a telomere-binding protein. Co-staining with **DST20** and TRF2 antibodies demonstrated clear nuclear colocalization of **DST20** and TRF2 signal in both U2OS and HeLa cells (Fig. 4b,c), with ∼20% of **DST20** signals overlapping TRF2 foci in U2OS cells (Fig. 4).

**Figure 4:**
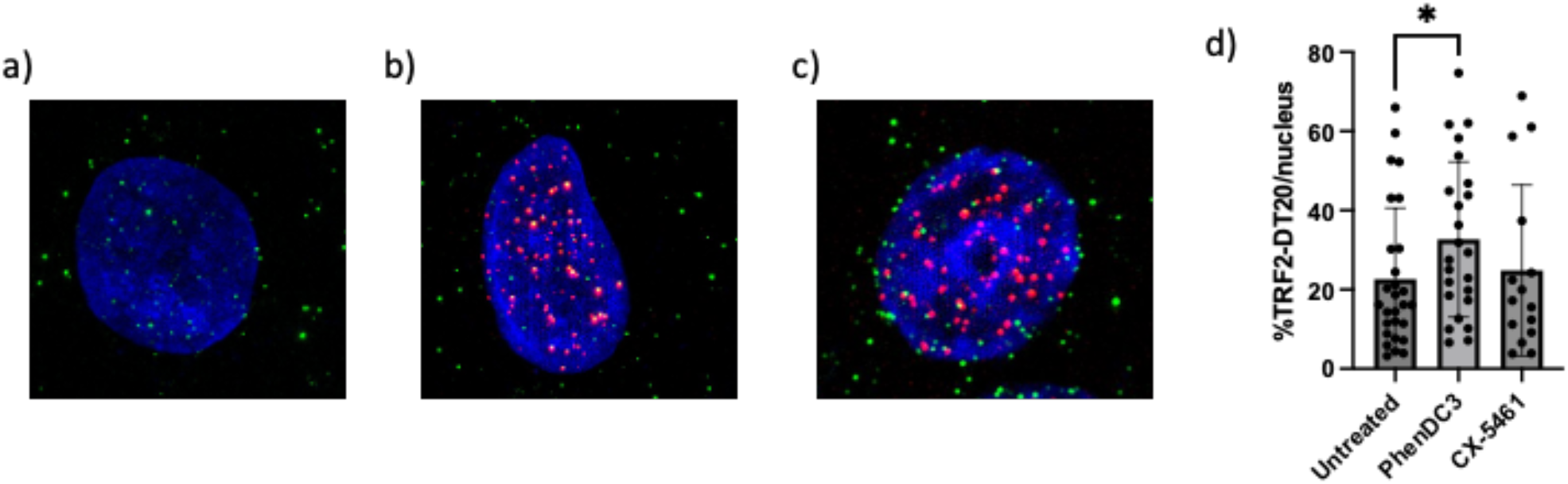
Visualization of telomeric G4 structures in human cancer cells. **a)** Immunofluorescence showing DST20 foci in U2OS cells per nuclei, **b)** Co-localization of TRF2 protein (red) and *IgG* **DST20** signals (green) in U2OS, **c)** Co-localization of TRF2 protein (red) and *IgG* **DST20** signals (green) HeLa cells, **d)** Quantification of TRF2 foci co-localizing with *IgG* **DST20** in untreated and PhenDC3 or CX-5461 treated U2OS cells. *Error bars represent SD from the means, *n* = 4 independent experiments. p values were calculated using unpaired Student’s *t* tests. ns: p>0.05; *p<0.05; **p<0.01; ***p<0.001.

To further investigate the selective binding of **DST20** to antiparallel G4 structures formed by telomeric sequences in cells, we hypothesized that stabilization of these structures by G4-binding ligands would increase the colocalization of **DST20** with TRF2 foci. U2OS cells were treated with PhenDC3, a well-characterized G4 ligand known to preferentially bind telomeric G4 structures, ^22^ and with CX-5461, a G4 ligand primarily targeting ribosomal DNA sequences.^23^ Upon treatment with PhenDC3, a significant increase in the number of TRF2 foci colocalizing with **DST20** signals was observed, whereas no such increase was detected following treatment with CX-5461 (Fig. 4d).

These results demonstrate that the *IgG* **DST20** antibody specifically recognizes a subset of genomic G-quadruplex structures, with a significant fraction localized at telomeres. Furthermore, the increased colocalization observed upon treatment with PhenDC3—but not with CX-5461—strongly suggests that *IgG* **DST20** preferentially binds to antiparallel G4 structures stabilized at telomeres.

## Conclusion

In summary, we have developed a novel highly specific DNA G-quadruplex antibody **DST20** and employed it to visualize G-quadruplex structures in the DNA of human cells. ELISA assays and BLI studies demonstrated that *IgG* **DST20** antibody is clearly selective for the antiparallel G4 topology of the telomeric sequence. Alternative antiparallel sequence such as 22CTA and hybrid sequence such as HIV-LTRIII are also not recognized by this antibody which suggest a crucial structural feature for the recognition. Likewise, the affinity of this new antibody for parallel G4 topology such as in CEB25, KIT and MYC sequences is much weaker thus allowing an efficient discrimination. Finally, *IgG* **DST20** does not appear to bind to the over abundant duplex DNA or RNA sequences.

At the cellular level, the partial colocalization of *IgG* **DST20** with the telomeric protein TRF2, combined with the enhanced overlap observed upon treatment with the telomere-targeting G4 ligand PhenDC3—but not with the rDNA-specific ligand CX-5461—supports the notion that this antibody preferentially binds to antiparallel G4 conformations stabilized at telomeres. Since *IgG* **DST20** preferentially recognizes G-quadruplex structures at telomeres, it serves as an ideal tool for investigating the role of these structures in the maintenance of telomeric sequences or studying the action of ligands at the telomere level. Also, in our point of view, *IgG* **DST20** antibody do not compete with the commercially available BG4 or 1H6 but rather should be associated with them to have a more precise view of the G-quadruplexes repartition in the cells.

## Acknowledgements

We thank the French National Research Agency (grants ANR-16-CE11-0006-01 and ANR-21-CE44-0005-02) and University Grenoble Alpes Graduate School (ANR-17-EURE-0003) for financial support. NanoBio-ICMG (UAR 2607) is acknowledged for their support for BLI studies. We acknowledge TRI-IPBS imaging facilities (France-BioImaging ANR-24-INBS-0005 FBI BIOGEN).

## Author contributions

N.R., S.M., C.E., Em.D. carried out the antibodies selection experiments, H.B. and Em.D. performed the BLI experiments, M.F.S and D.G. carried out experiments on cells, E.P. performed image analyses quantifications, T.L synthetized the molecular systems **1** and **2**, and J.D., N.S. and E.D. designed the experiments. T.L., D.G., J.D., N.S. and E.D. co-wrote the manuscript.

## Competing interests

The authors declare no competing financial interests.

## Additional information

Supplementary information is available in the online version of the paper. Reprints and permission information is available online at http://www.nature.com/reprints. Correspondence and requests for materials should be addressed to E.D.

## Notes

### Competing Interest Statement

The authors have declared no competing interest.

