## Supporting Information for "Antibody targeting the anti-parallel topology of human telomeric G-quadruplex DNA"

**Summary**

**Methods**

Phage display selection………………………………………………………………………...	p2

*IgG* production……………………………………………………………...............................	p2

Enzyme-linked immunosorbent assay (ELISA)………………………………………….….	p2

Bio Layer Interferometry (BLI)………………………………………………………………...	p3

Cell cultures and immunofluorescence……………………………………………………….	p3

**Enzyme-linked immunosorbent assay (ELISA)**

**Figure S1**: Screening by Phage ELISA………………………………………………………	p5

**Figure S2**: Comparison with BG4…………………………………………………………….	p5

**Figure S3**: Binding curves for the interaction of scFv **DST20** with various DNA and RNA targets……………………………………………………………………………………………	p6

**CD analysis**

**Figures S4-S6**: Spectra of the various oligonucleotides……………………..……………p7-8

**BLI studies**

**Figure S7**: Incubation of *IgG* **DST20** at 2 nM with different DNA et RNA sequences.….	p9

**Figure S8**: BLI analysis of binding of different concentrations of **DST 20** ………………	p10

**Cellular studies**

**Figure S9**: Fluorescence microscopy image acquired in the absence of DST20 treatment………………………………………………………………………………………………	p11

**Phage display selection**

The single-chain variable fragment (*scFv*), **DST20**, was isolated from the Tomlinson I library (1,47× 10^8^ *scFv* clones) through selection using the constrained antiparallel G4 **2**. Phage production, phage display screens based on the pIT2 phagemid vector, and helper phage KO7 production were performed following the standard protocol from our own previous work.^[1]^

The DNA target **2** was resuspended in Tris-EDTA buffer at 400 µM. For each selection round, 50 µL of magnetic beads coated with streptavidin (Dynabeads(R) M-280 Streptavidin from Invitrogen Life Technologies SAS, Saint Aubin, France) were prepared according to the manufacturer’s protocol. 10 µL of DNA stock solution were mixed with 50 µL of washed Dynabeads (R) and incubated for 10 minutes at room temperature using gentle rotation. The biotinylated DNA coated beads were separated with a strong magnet for 2-3 minutes and washed 2-3 times with a buffer containing 5 mM Tris-HCl (pH 7.5), 0.5 mM EDTA and 1M NaCl. Phages were first incubated against either naked magnetic beads for negative selection. For DNA target selection, phages were incubated during 1 h without agitation and 30 minutes on a rocker at room temperature and 10 washing steps with 1xPBS + 0.1% Tween 20 were performed.

***IgG* production**

The selected *scFvs* was reformatted into *IgG* format (with a human IgG1/κ isotype for both antibodies) by transient transfection into CHO cells, and produced in PBS buffer (pH 7,4; NaCl 137mM - KCl 2,7mM - Na_2_HPO_4_ 10mM - KH_2_PO_4_ 2mM). *IgG* **DST20** were produced by BIOTEM.

**Enzyme-linked immunosorbent assay (ELISA)**

ELISA assays for binding affinity and specificity were performed using standard methods. Briefly, biotinylated oligonucleotides were bound to streptavidin-coated plates (Thermo-scientific-43601-Nunc immobilizer streptavidin F96 clear-batch 167442) followed by incubation with **DST20**, and detection was achieved with an Anti-Human IgG1-peroxidase antibody (Mouse monoclonal, SAB4200768-1VL) and TMB (Tetramethylbenzidine,Thermo Scientific-TMB substrate kit-34021, batch WF333257). Signal intensity was measured at 450 nm on a TECAN microplate reader (infinite M200 PRO). Dissociation constants (*K*_D_) were calculated from fitted binding curves using Origin software (^©^OriginLab Corporation) and error bars represent standard error means calculated from 3 replicates. The different oligonucleotides were annealed in PBS, pH 7.4, by slow cooling from 95 °C to room temperature.

**Bio Layer Interferometry (BLI)**

*Experiment***.** The streptavidin biosensors (Octet^®^ SA Biosensors, batch 2107002711, 18-0019) were functionalized with 100 nM biotinylated oligonucleotides. The immobilization time depends on the experiments and were adjusted to obtained the better recognition results. Biosensors are then immersed in wells containing **DST20** antibody at different concentrations (association step) and back to the buffer (dissociation step). During all these steps, the changes in the wavelength of the reflected light coming from the surface of the biosensors are registered by the Octet^®^ BLI system. Reference sensors without DNA immobilization were used to subtract the non-specific adsorption on the SA layer. One sensor was used *per* concentration due to the weak dissociation and to avoid the use of regenerative solution. G-quadruplex oligonucleotides were annealed in 10 mM PBS, pH 7.4, by slow cooling from 95 °C to room temperature.

*Data analysis.* All association steps were fitted using a 1:1 model to determine the kinetic association rate *k_on_* and dissociation rate *k_off_*. The dissociation and association equilibrium constants (*K*_D_ and *K*_A_, respectively) were calculated from the binding rate constants as *K_D_* = *k*_off_/*k*_on_ or *K*_A_ = *k*_on_/*k*_off_ or determined by the fitting of the Langmuir isotherm from the response at the equilibrium state. The reported values were obtained from the average of independent experiments, and the errors provided are standard deviations relatively to the mean. Each experiment was repeated at least two times.

**Cell cultures and immunofluorescence.**

All the cell lines used in this study were obtained from ATCC that certified their identity. DAPI staining analysis was used to confirm the absence of *Mycoplasma* contamination during the course of experiments. All culture media were provided by Gibco and were supplemented with 10 % fetal bovine serum (Eurobio), 100 U/mL penicillin (Gibco), and 100 µg/mL streptomycin (Gibco). Cells were grown in a humidified atmosphere with 5 % CO_2_ at 37 °C. HeLa and U2OS cells were grown in Dulbecco’s Modified Eagle Medium.

HeLa and U2OS cells were seeded in 24-wells plate at 100,000 cells/well on #1.5 glass coverslips (VWR, #631-0150). 24 h later, cells were treated with PhenDC3 and CX-5461 used respectively at 20 μM and 0.2 µM for 8 h. Then, cells were washed with ice-cold PBS and then incubated twice for 3 minutes at room temperature with CSK buffer (10 mM Pipes, pH 7.0, 100 mM NaCl, 300 mM sucrose, and 3 mM MgCl_2_) containing 0.7% Triton X-100 (CSK) and 0.3 mg/ml RNase A. After pre-extraction, cells were washed with PBS and fixed with 2% PFA for 20 minutes. For immunodetection cells were washed with PBS and incubated for about 1 h at room temperature in blocking buffer (20 mM Tris-HCl pH 7.5, 150 mM NaCl, 2% BSA, 0.2% fish gelatin, 0.1% Triton-X 100) prior to incubation overnight at 4 °C with primary antibody diluted in blocking buffer. Cells were then washed with PBS 0.1% Tween-20 and incubated with appropriate secondary goat antibody coupled to Alexa Fluor 488 or 594 diluted in blocking buffer (for 1 h at room temperature. At last, cells were washed with PBS 0.1% Tween-20 and stained with 0.1 μg/mL DAPI for 20 minutes at room temperature, and coverslips were mounted with Vectashield mounting medium (Vector Laboratories). The **DST20** antibody was used at a concentration of 2 µg/mL. For telomeric staining, anti-TRF2 (clone 4A794, Abcam) was used at a 1:1,000 dilution. Alexa Fluor 488-conjugated anti-human (A11013) and Alexa Fluor 594-conjugated anti-mouse secondary antibodies (A11012) (Thermo Fisher Scientific) were both used at 1:1,000 dilution. Imaging was performed using a Zeiss Elyra 7 3D Lattice SIM super-resolution microscope equipped with a 40× PLAN APO NA 1.4 objective and dual sCMOS cameras (pco.edge). 3D-SIM image reconstructions were carried out using Zen Black 2.3 software (Zeiss).

Image analysis was performed using Imaris software (RRID:SCR_007370). Nuclear segmentation was carried out manually using the Surface module. The 3D localization of antibody signals was analyzed using the Spot detection function, with the spot diameter set to 0.4 µm, corresponding to the average measured size of individual foci. Spots were classified as either intra-nuclear or external relative to the nuclear surface using the "Spots to Surfaces" tool, with the distance threshold set to 0 µm. Co-localization analysis between spots was performed using the "Spots to Spots" function in Imaris, with a maximum distance threshold equal to the spot diameter (0.4 µm), thereby defining co-localization as direct contact or overlap between spots.

**Enzyme-linked immunosorbent assay (ELISA)**

**Screening by Phage ELISA**

The tests were conducted on the supernatants from bacterial cultures infected with each phage, after induction with IPTG. This process releases *scFv* fragments fused to the PIII protein of phage M13, as described in the protocol.

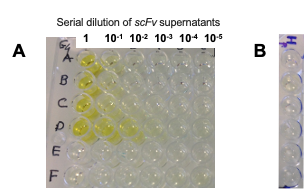

**Figure S1.** Example of assays performed on approximately one hundred *scFvs* selected by phage display. The quantity of *scFv* produced was assessed by ELISA against the telomeric G4 **2** target (panel **A**), using a range of *scFv* concentrations. For the HP-DNA target (panel **B**), only the highest *scFv* concentration was tested. **DST20** is shown in row D, which corresponds to the dilution (10⁻³) where a positive signal remains detectable. All *scFvs* tested in this assay were negative against HP-DNA.

**Comparison with BG4**

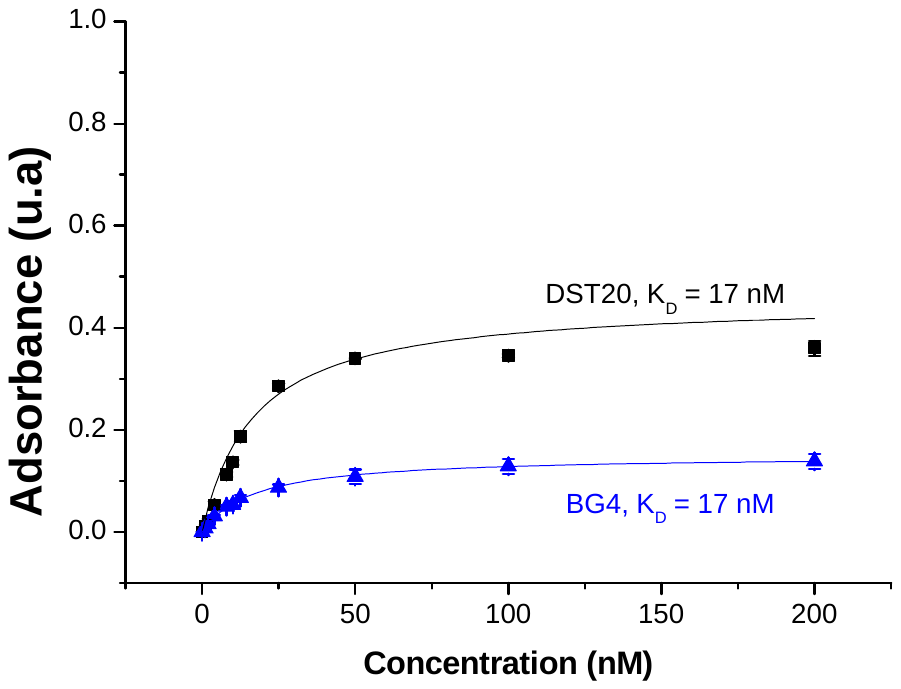

**Figure S2.** Binding curves as determined by ELISA assay for the interaction of *IgG* BG4 and *IgG* **DST20** with Tel23.

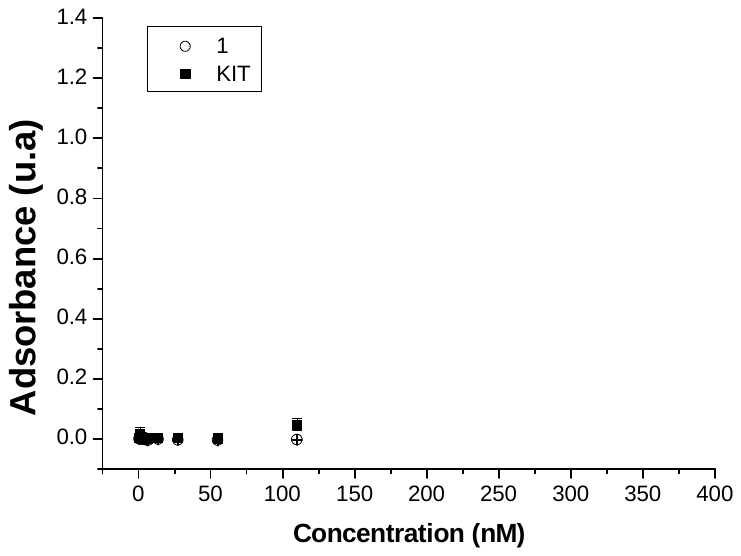

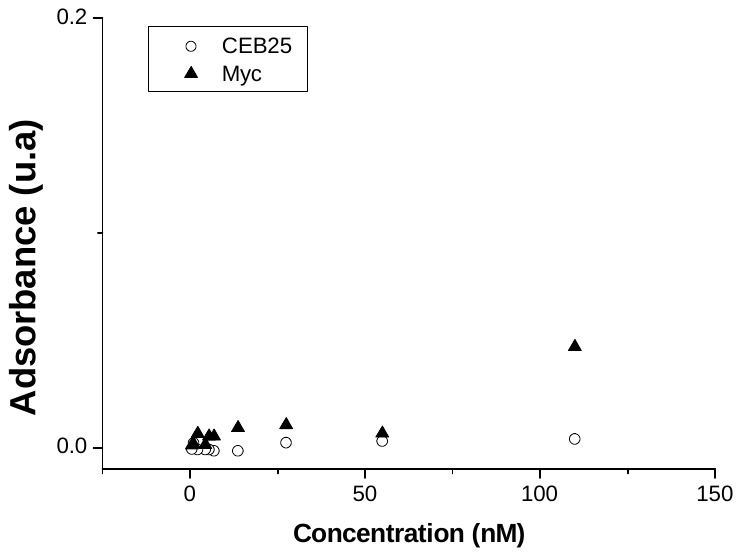

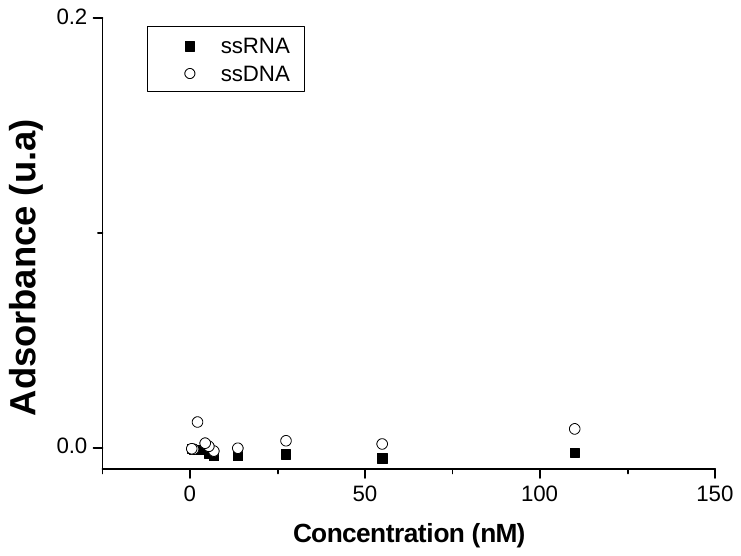

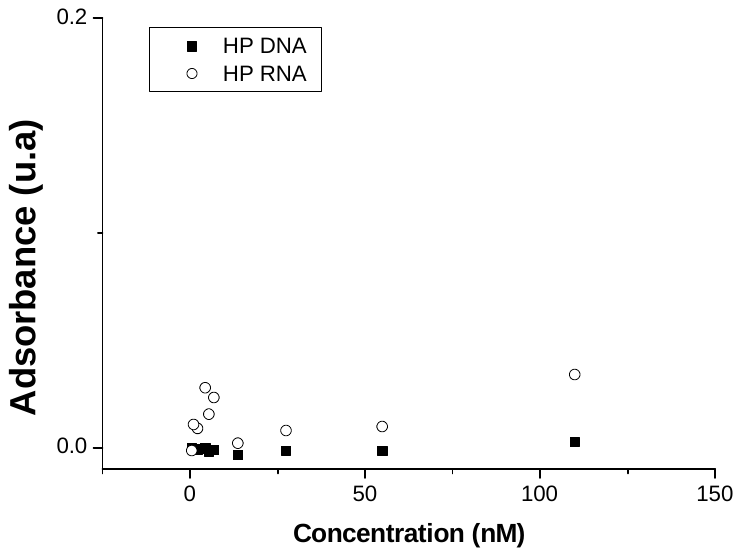

**a)**

**b)**

**c)**

**d)**

**e)**

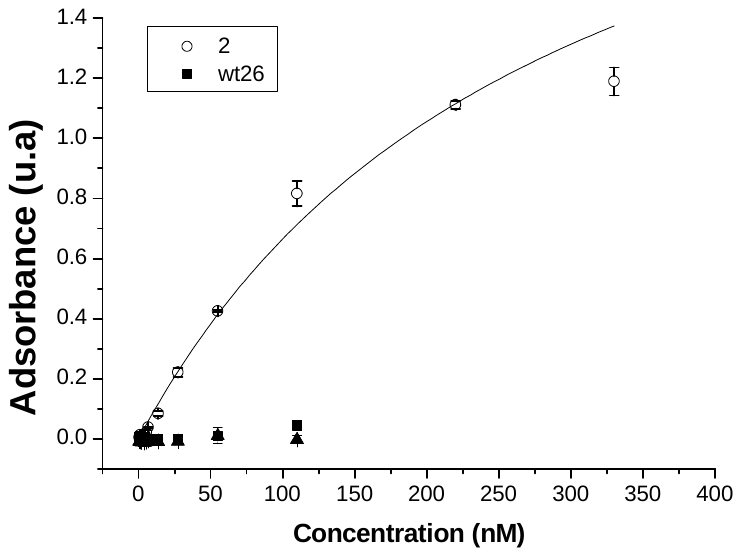

**Figure S3:** Structure specificity of the *scFv* **DST20** antibody for G-quadruplex structures. Binding curves as determined by ELISA with **a)** Constrained antiparallel G-quadruplex **2** and mixture of parallel and antiparallel topologies wt26, **b)** parallel G-quadruplexes KIT and constrained structure **1**, **c)** parallel G-quadruplexes MYC and CEB25, **d)** double stranded DNA and RNA and **e)** single stranded DNA and RNA.

**CD analysis**

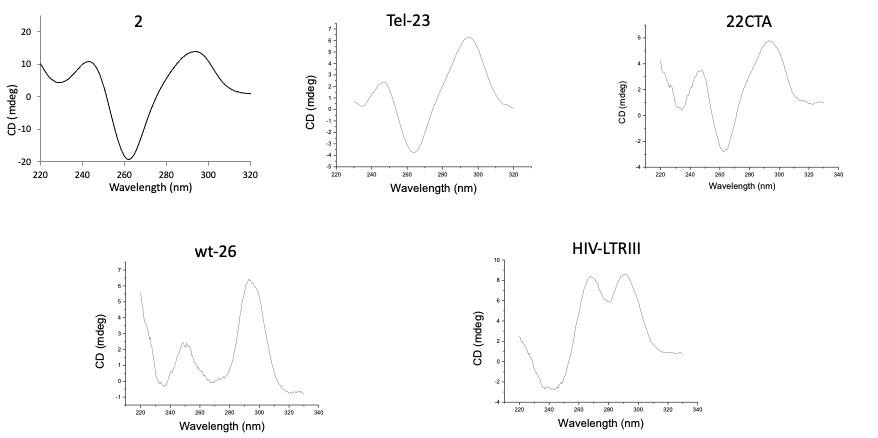

**Figure S4.** CD spectra of G-quadruplex oligonucleotides for selectivity studies. CD analyses were carried out in PBS buffer at pH 7.2 in the same conditions used for phage display selection.

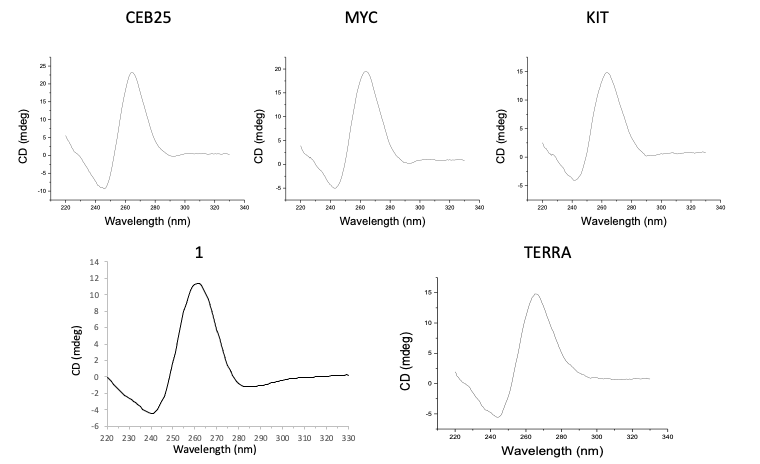

**Figure S5.** CD spectra of the parallel G-quadruplex oligonucleotides for selectivity studies. CD analyses were carried out in PBS buffer at pH 7.2 in the same conditions used for phage display selection.

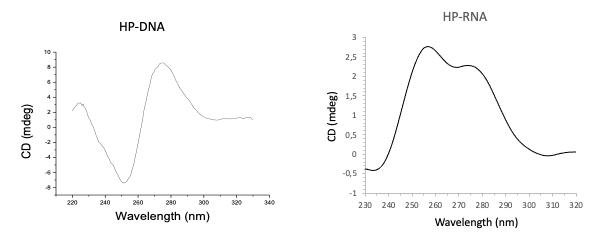

**Figure S6.** CD spectra of the hairpin oligonucleotides for selectivity studies. CD analyses were carried out in PBS buffer at pH 7.2 in the same conditions used for phage display selection.

**BLI studies**

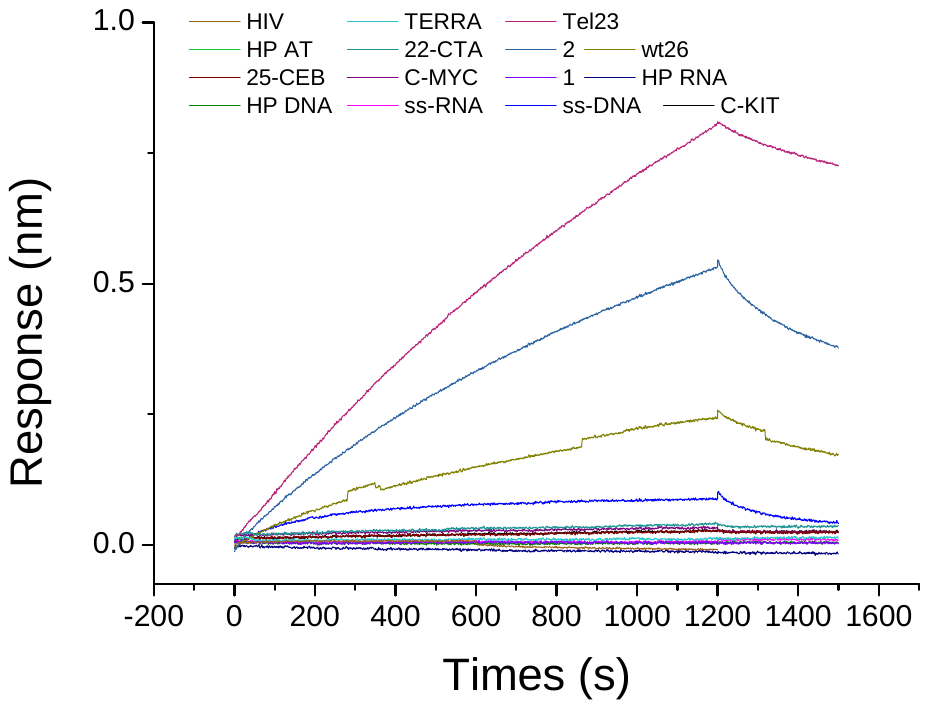

Tel23

Constrained system **2**

wt26

ssDNA

**Figure S7**. Sensorgrams recorded during the incubation of *IgG* **DST20** at 2 nM with different DNA et RNA sequences for 20 minutes.

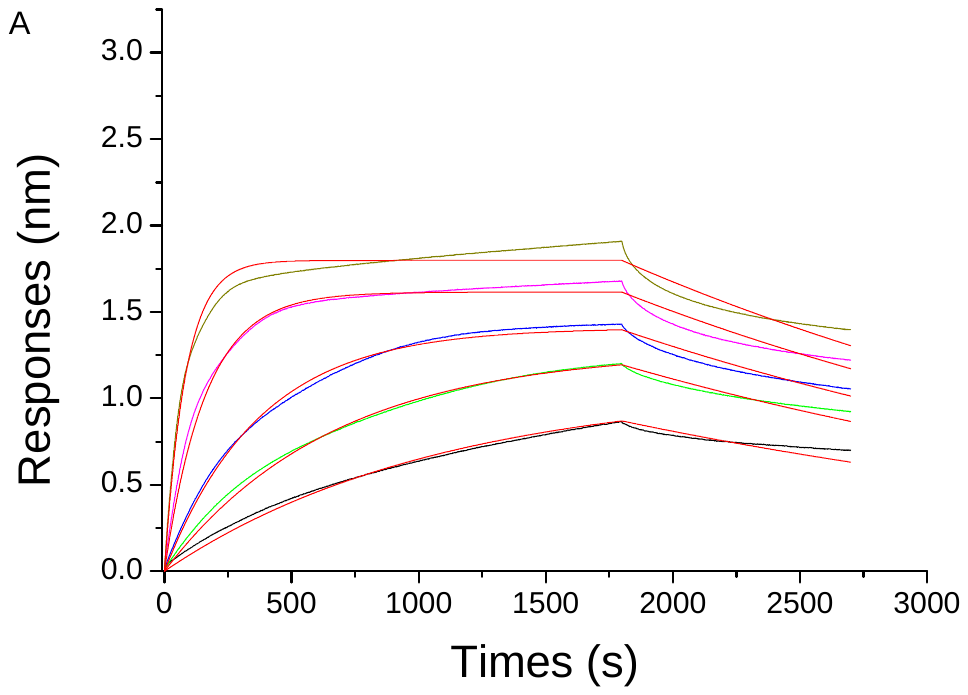

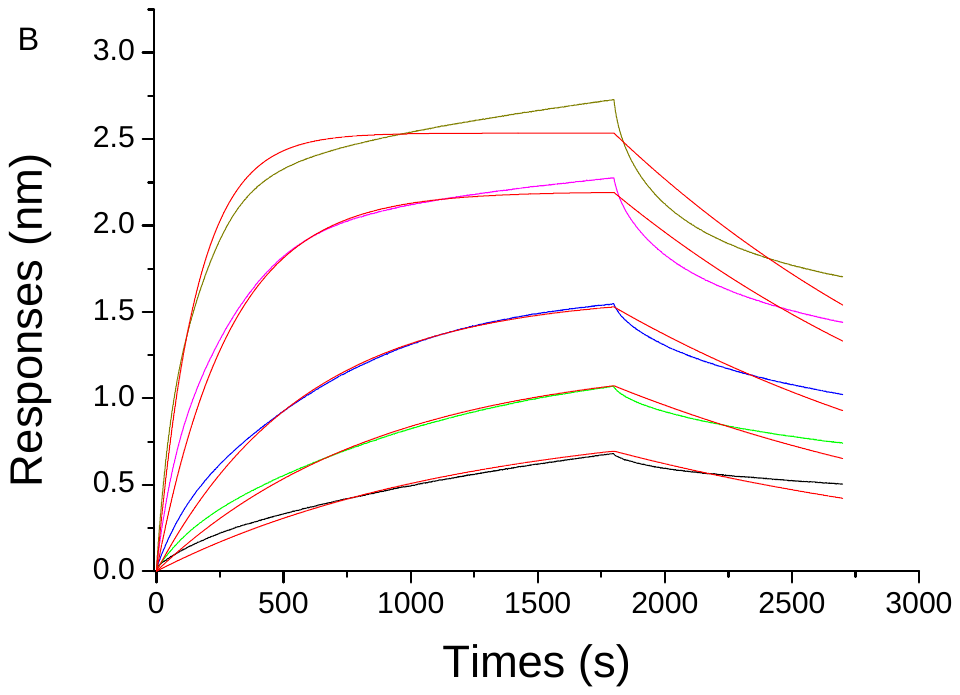

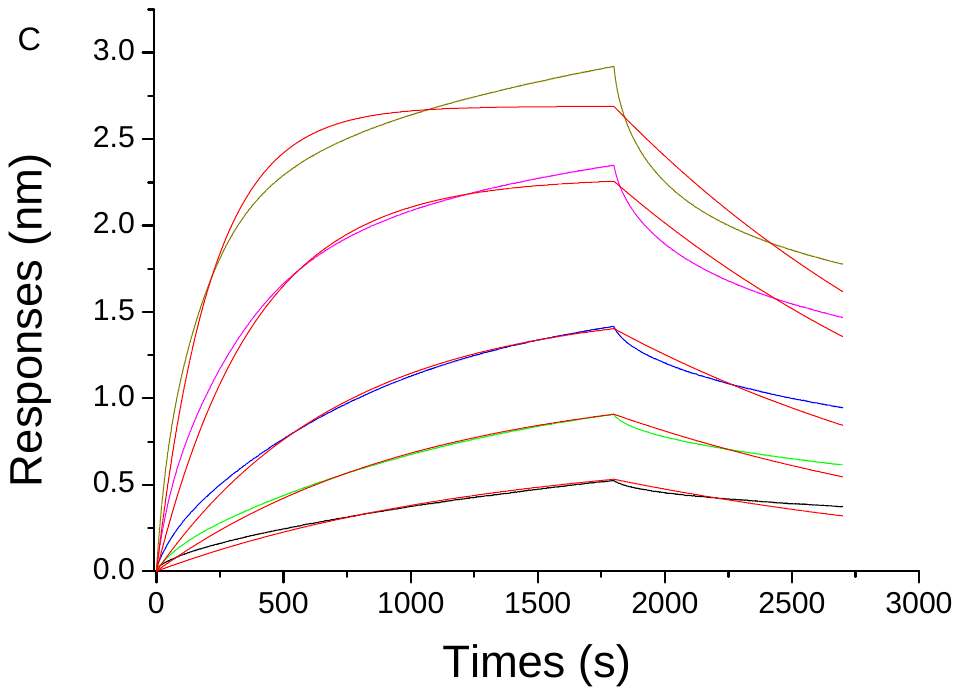

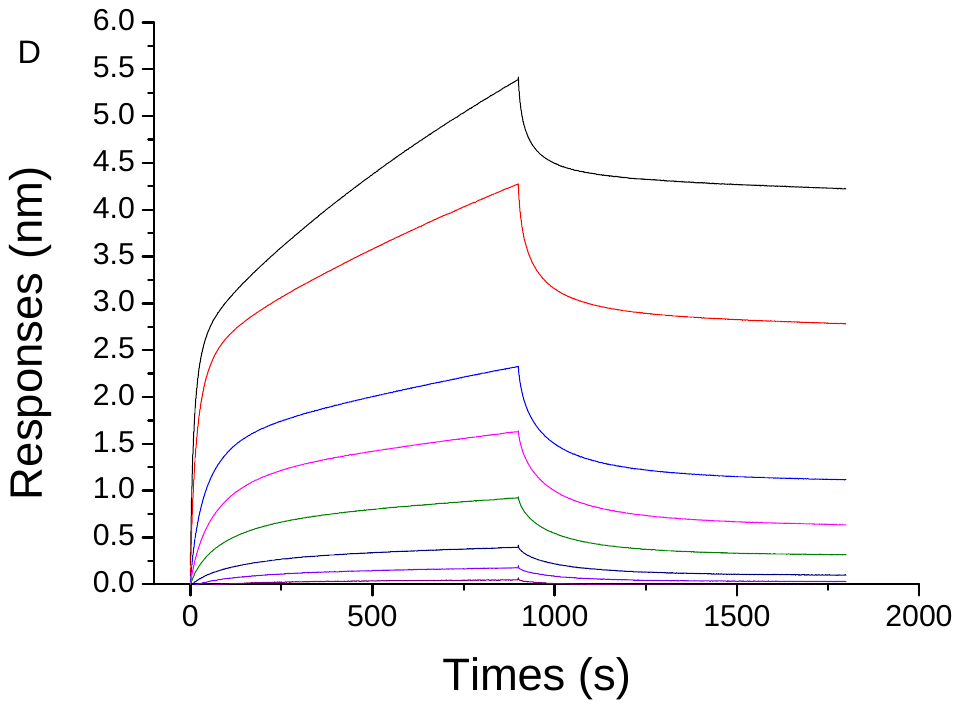

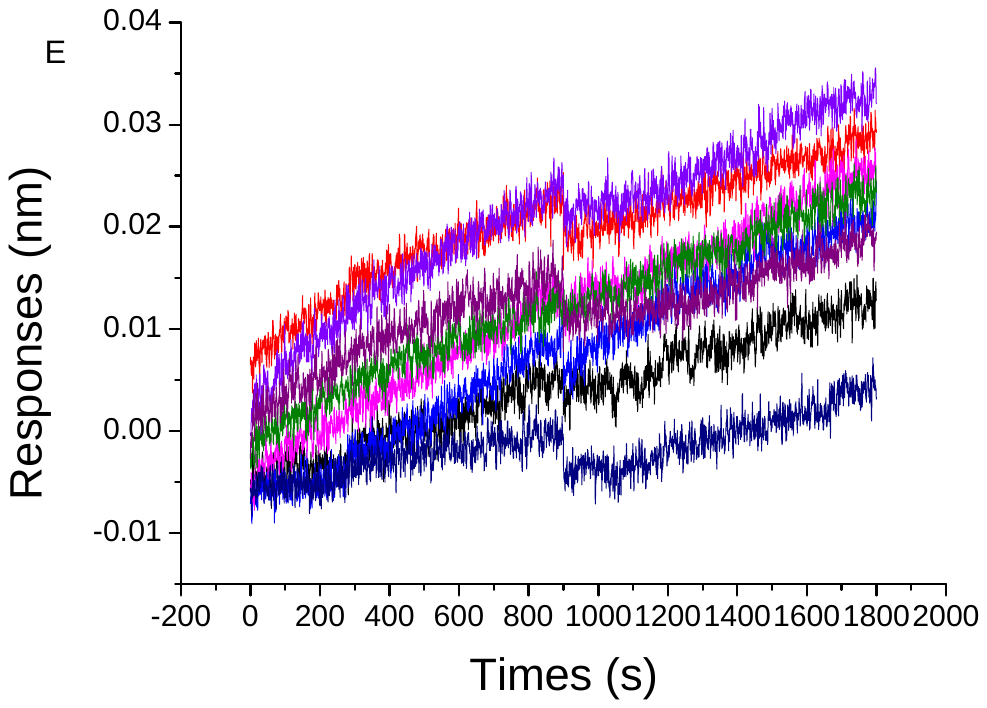

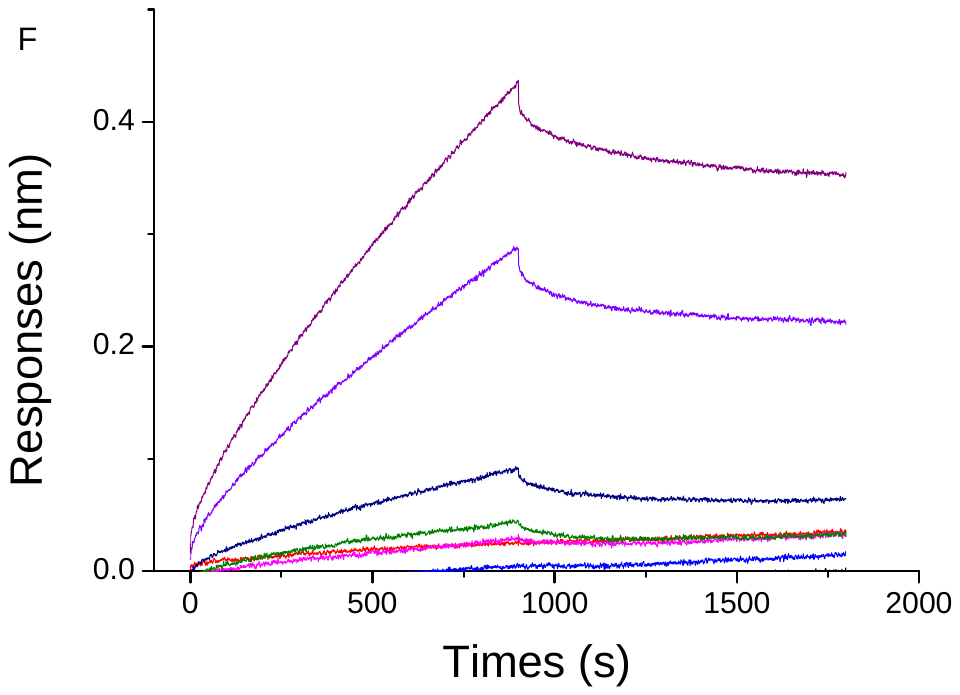

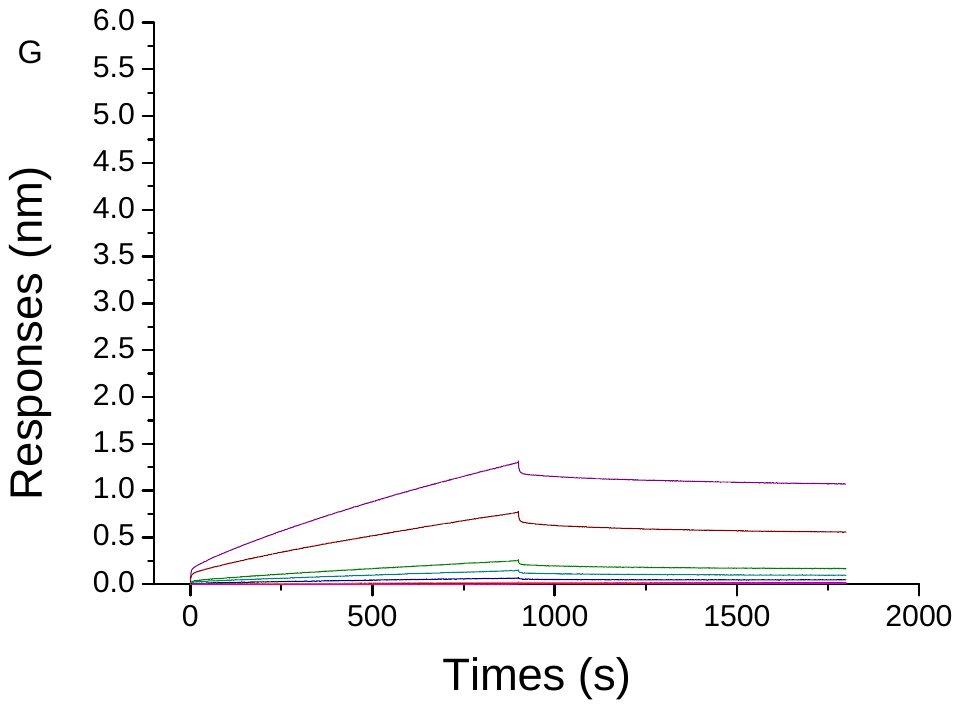

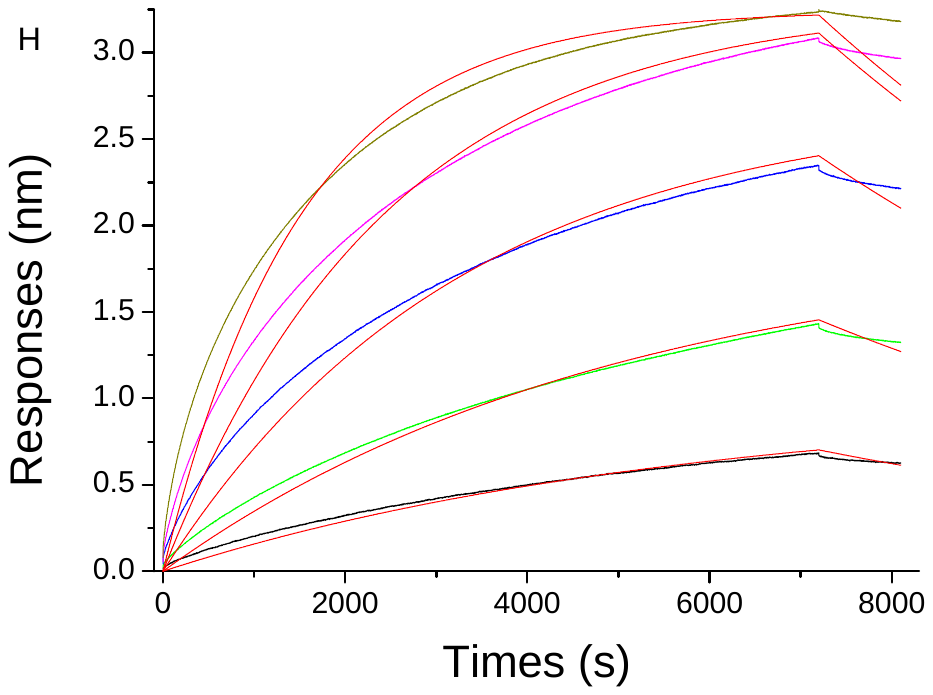

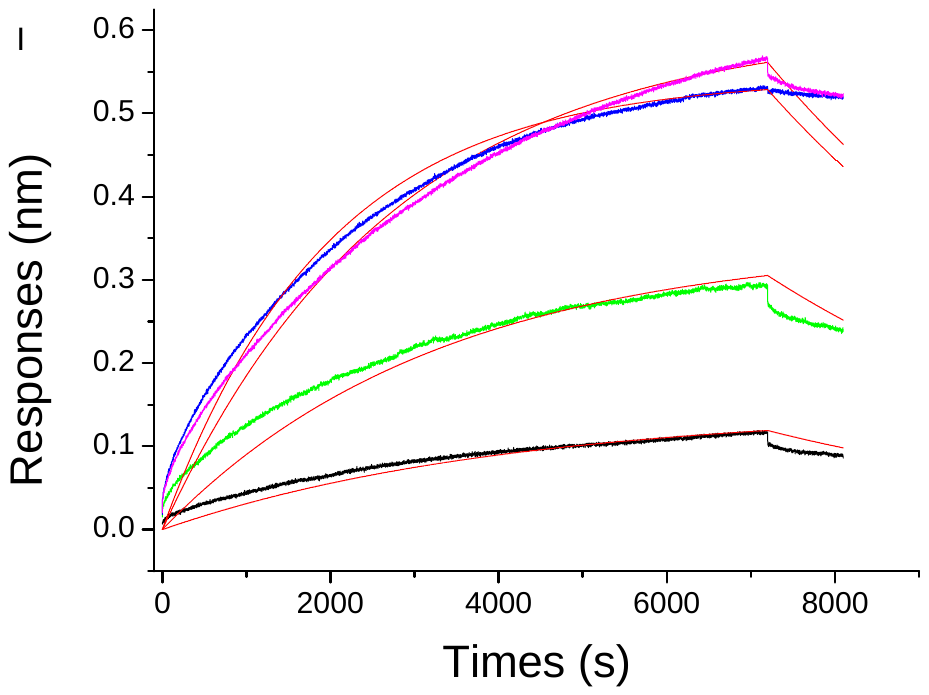

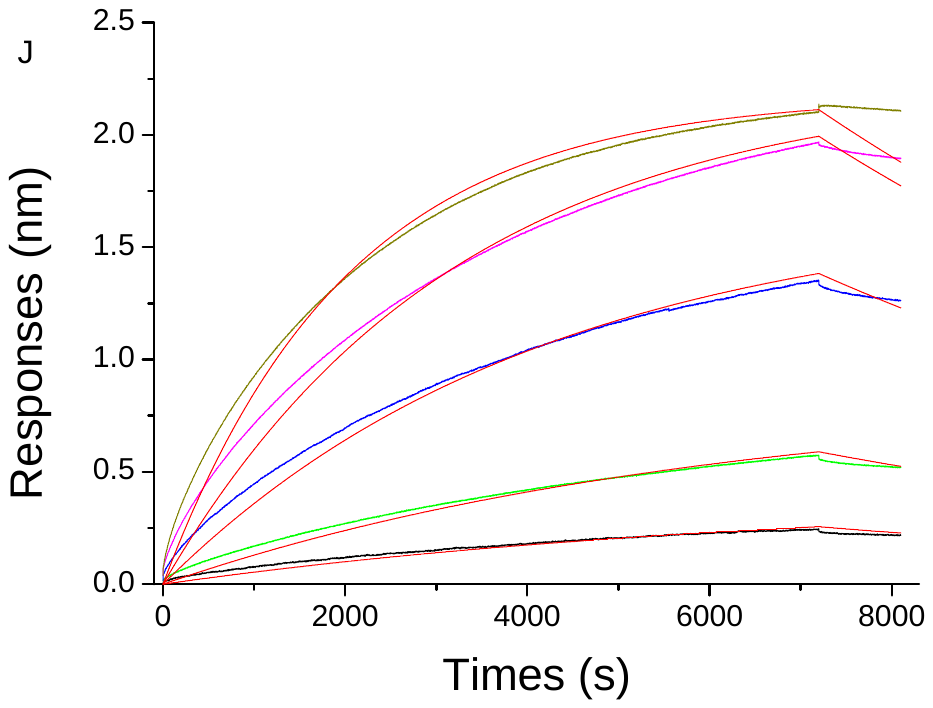

**Figure S8.** BLI analysis of binding of different concentrations of **DST 20** to **2** (**A**), Tel-23 (**B**), wt26 (**C**), ssDNA (**D**), ssRNA (**E**), HP DNA (**F**), 22CTA (**G**), CEB-25 (**H**), MYC (**I**) and KIT (**J**). Color lines (excepted red) are experimental curves; red lines are the fits to the 1:1 model. The concentration range was 2.5, 5, 10, 25 and 50 nM for **A-C**, 2.5, 5, 10, 25, 50, 100, 300 et 500 nM for **D-G** and 50, 100, 300, 500, and 1000 nM for **H-J**.

**Cellular studies**

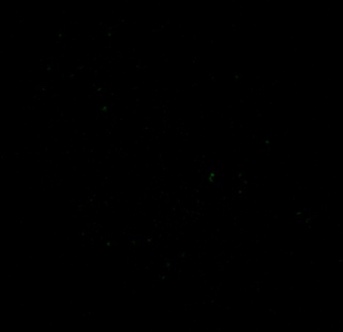

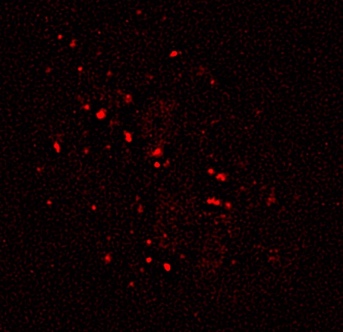

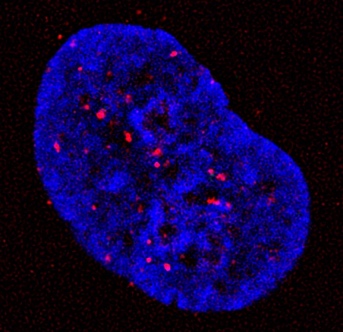

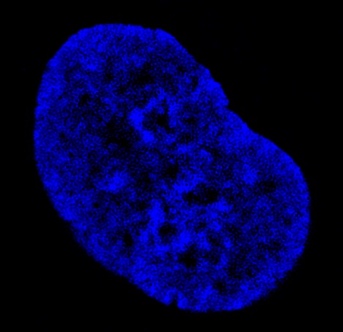

**Figure S9.** Fluorescence microscopy image acquired in the absence of DST20 treatment. From left to right: TRF2 signal (red channel), Alexa488 signal in the absence of DST20 (green channel), nuclear counterstaining with DAPI (blue channel), and merged image showing the overlay of all three channels.

1. Boyer, S., Biswas, D., Kumar Soshee, A., Scaramozzino, N., Nizak, C., Rivoire, O. Hierarchy and extremes in selections from pools of randomized proteins, *Proc. Natl. Acad. Sci. U.S.A.* **113**, 3482-3487, (2016).
